# An Omni-Mesoscope for multiscale high-throughput quantitative phase imaging of cellular dynamics and high-content molecular characterization

**DOI:** 10.1101/2024.07.18.604137

**Authors:** Hongqiang Ma, Maomao Chen, Jianquan Xu, Yongxin Zhao, Yang Liu

## Abstract

The mesoscope has emerged as a powerful imaging tool in biomedical research, yet its high cost and low resolution have limited its broader application. Here, we introduce the Omni-Mesoscope, a cost-effective high-spatial-temporal, multimodal, and multiplex mesoscopic imaging platform built from cost-efficient off-the-shelf components. This system uniquely merges the capabilities of quantitative phase microscopy to capture live-cell dynamics over a large cell population with highly multiplexed fluorescence imaging for comprehensive molecular characterization. This integration facilitates simultaneous tracking of live-cell morphodynamics across thousands of cells, alongside high-content molecular analysis at the single-cell level. Furthermore, the Omni-Mesoscope offers a mesoscale field of view of approximately 5 mm^2^ with a high spatial resolution down to 700 nm, enabling the capture of information-rich images with detailed sub-cellular features. We demonstrate such capability in delineating molecular characteristics underlying rare dynamic cellular phenomena, such as cancer cell responses to chemotherapy and the emergence of polyploidy in drug-resistant cells. Moreover, the cost-effectiveness and the simplicity of our Omni-Mesoscope democratizes mesoscopic imaging, making it accessible across diverse biomedical research fields. To further demonstrate its versatility, we integrate expansion microscopy to enhance 3D volumetric super-resolution imaging of thicker tissues, opening new avenues for biological exploration at unprecedented scales and resolutions.

## Introduction

Biological systems are characterized by significant cellular and molecular heterogeneity, functioning within diverse microenvironments across various time scales (*1, 2*). Addressing this complexity requires technologies capable of simultaneously capturing both the behaviors and molecular characteristics of individual cells over time across a large cell population in their spatial context at a high temporal and spatial resolution. Such a high-throughput high-resolution imaging system is crucial for identifying dynamic, rare events coupled with detailed sub-cellular and molecular characteristics, thereby facilitating in-depth analysis of diverse cellular phenotypes. This multiscale multi-functional imaging approach is crucial in revealing the mechanisms underlying therapeutic response from a systems-to-molecular perspective and in identifying effective diagnostic or prognostic biomarkers.

Traditional imaging systems are often limited by a compromise between the field of view and image resolution, significantly hindering our ability to simultaneously observe cellular dynamics with the necessary sub-cellular details across a large cell population. The emergence of mesoscopic imaging technology represents a significant advancement, offering an ultra-large field of view spanning several millimeters (*3*). However, most mesoscopic systems are limited to a resolution of several microns (*4*–*10*) due to various factors such as numerical aperture (NA), optical aberrations, magnification, and image sensors. Although existing mesoscope allows us to observe cell dynamics over a large cell population, their limited resolution hampers the visualization of the subcellular details. To enhance spatial resolution, specially designed mesoscopic objective lenses with a higher NA (up to 0.47) (e.g., mesolens (*11*)) have been developed, but their widespread adoption is hindered by the complexity and cost of manufacturing. Moreover, conventional mesoscope often relies on a few fluorescent markers, which can perturb the natural state of cells, thus limiting the capacity for long-term observation of live-cell dynamics with rich functional insights. These challenges underscore the urgent need for an advanced multimodal imaging solution that can overcome these limitations to observe cell dynamics at a high spatiotemporal resolution over a large cell population without sacrificing in-depth functional information.

We present the Omni-Mesoscope, a versatile and cost-effective multi-modal mesoscopic imaging system that achieves a sub-micron spatial resolution across a wide field of view, approximately 5mm^2^, at a speed of 4 frames per second, primarily limited by the camera speed. Importantly, Omni-Mesoscope couples quantitative phase microscopy for label-free live-cell imaging with highly multiplex fluorescence microscopy for high-content functional imaging.

This integration allows for continuous monitoring of live-cell dynamics alongside detailed molecular analysis on the same cells over a large cell population, providing a comprehensive view of large-scale dynamic cellular processes and their underlying molecular characteristics. We demonstrate the capability of Omni-Mesoscope to identify rare cell behaviors in response to chemotherapy drugs, which are linked to their underlying molecular characteristics. Furthermore, by incorporating expansion microscopy, we demonstrate the potential of Omni-Mesoscope for 3D volumetric super-resolution imaging of thick tissue across the mesoscale. This imaging system not only overcomes the limitations of conventional techniques but also opens new avenues for exploring the complexities of biological systems at unprecedented scales and resolutions.

## RESULTS

### Configuration and Performance Evaluation of the Omni-Mesoscope

Our Omni-Mesoscope is an automated imaging system with two major imaging modalities: label-free quantitative phase imaging module implemented via transport of intensity equation (TIE)-based phase retrieval; and highly multiplex fluorescence imaging module implemented with a flat-field illumination and multispectral detection. The schematic of the Omni-Mesoscope is shown in **Fig. 1**. Our system achieves an ultra-large field of view (FOV) at a high resolution through three key hardware elements. First, we adopted an off-the-shelf objective lens (MVPLAPO 2 XC, Olympus) at an NA of 0.5, originally designed for a stereo microscope.

**Figure 1.**
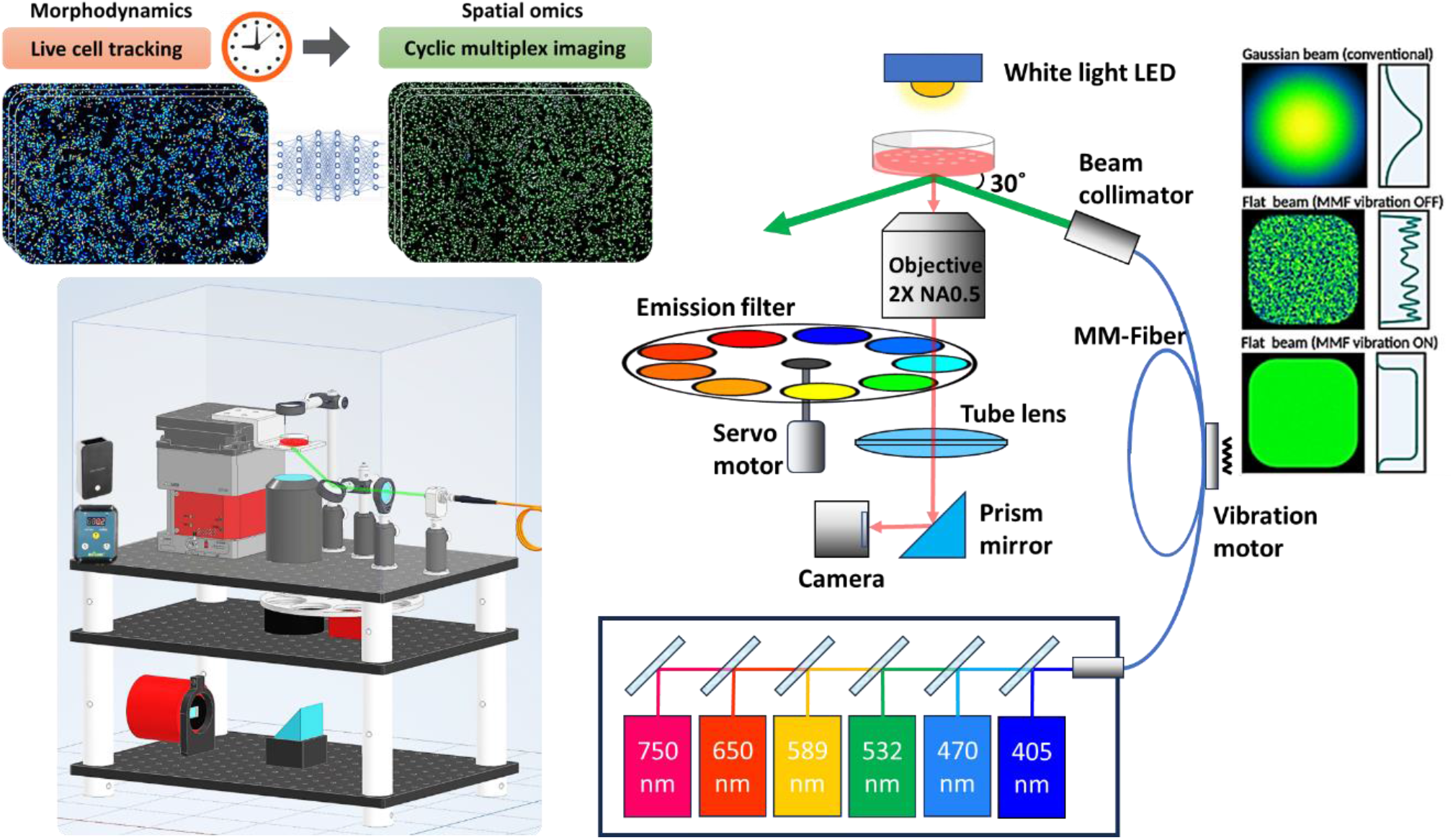
Schematics of our Omni-Mesoscope system. A collimated LED is used as the illumination beam for quantitative phase imaging. Six diode lasers (405nm, 470nm, 530nm, 589nm, 650nm, and 750nm), coupled to a multi-mode fiber and a high-frequency vibration motor to create uniform flat-field illumination. The illumination light was obliquely introduced from the bottom of the sample with an incident angle of ∼60°, which decouples from the detection path. A customized motorized filter wheel with 9 band-pass emission filters is used for multiplex fluorescence imaging.

Second, the mesoscope requires a large-format image sensor with tens of megapixels and a small pixel size, implemented with a cost-effective astronomy camera (ASI294MM pro, ZWO) that has a large-format sensor (IMX 492, ∼50 Megapixels, 12 times more than sCMOS) and a small pixel size of ∼2.3 μm. Third, a macro lens (DCR-5320 Pro, Raynox) generally used for photography was adapted as the tube lens to achieve an optical magnification of ∼7. This cost-efficient combination (∼$4,000) provides a final pixel size of ∼320 nm with a FOV of ∼5mm^2^ (2.8 mm ×1.8 mm).

A series of images of the fluorescent microspheres are captured along the axial dimension with a step size of 1μm. Then, we localized each from the three-dimensional fluorescent images, and the fluorescence intensity distribution of each microsphere was used to determine the point spread function (PSF) of the imaging system across the field of view. As shown in **Fig. 2**, the Omni-Mesocope achieves a lateral sub-micron resolution (700-900 nm) and an axial resolution of ∼6.2-8.7 μm across the entire field of view (2.8mm × 1.8mm). At the central region of the field of view, our system maintains a lateral resolution of about 700 nm. **Figure 2C** shows the field curvature of our system. Initially, the focus shift at the edges of the FOV can reach 8-10 μm; however, this is significantly reduced to less than 1μm after applying region-based post-refocusing (**Supplementary Figure S2**).

**Figure 2.**
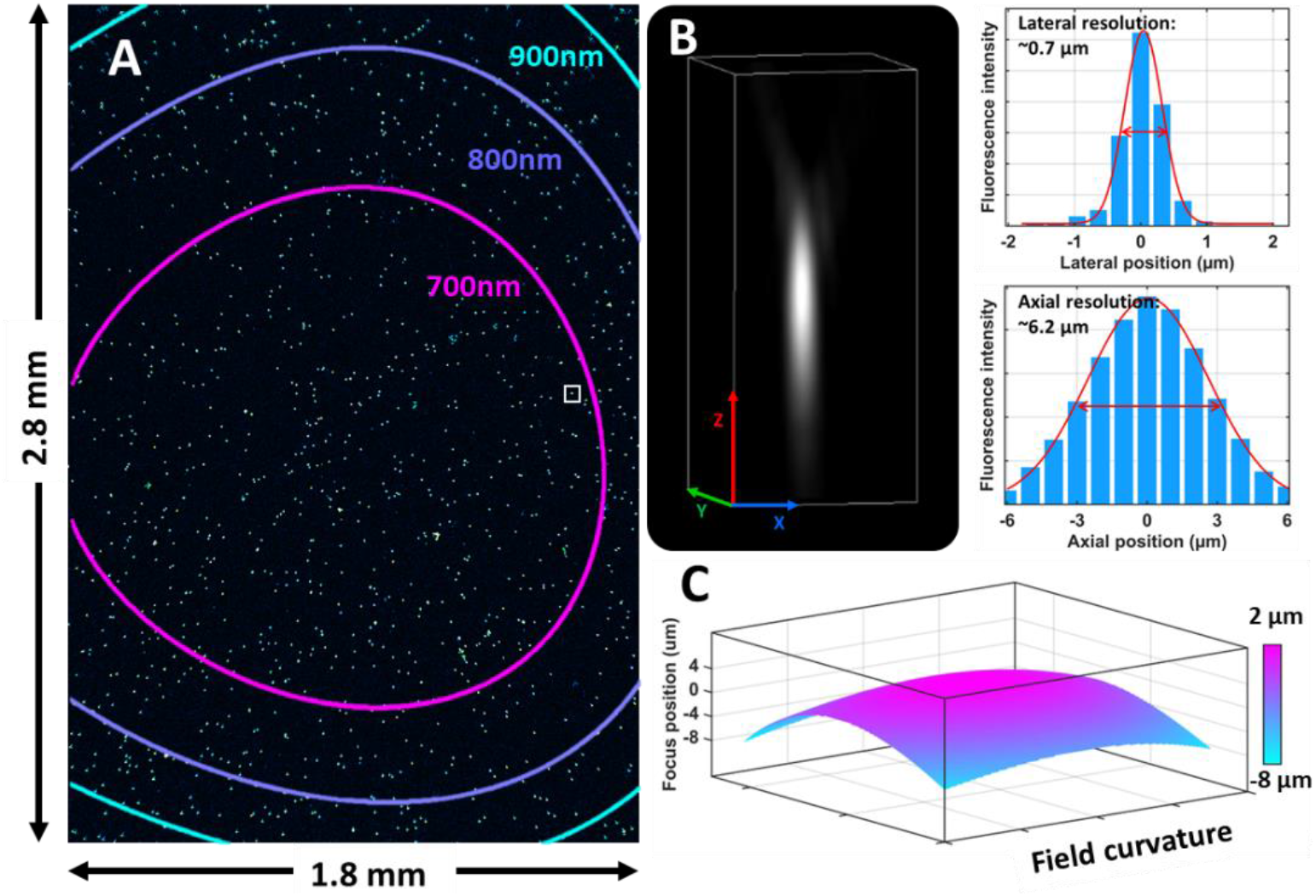
Performance evaluation of Omni-Mesoscope in resolution and field curvature. **(A)** The diffraction-limited image of fluorescent microspheres (FluoSpheres™, 0.2μm, 505/515) attached to the surface of a coverslip. To generate a precise and continuous map in resolution, we calculated the resolution at each position by averaging the resolution of tens of beads within a radius of 50 μm. **(B)** The point spread function (PSF) of the lateral and axial resolution. A Gaussian function is used to fit the PSF of each fluorescent microsphere and the full width at half maximum of the PSF is defined as the spatial resolution. **(C)** The field of curvature for Omni-Mesoscope. The focus shift at each position was calculated by averaging the axial position of all beads within a lateral radius of 50 μm.

The quantitative phase images are reconstructed based on the TIE-based phase retrieval (*12*) using standard bright-field intensity measurement at two different focal planes (±2 μm), which is simple and fast. In addition, our method does not require to modify the existing optical setup for fluorescence imaging compared to the interferometry-based quantitative phase imaging method (*13*–*16*). The accuracy of the quantitative phase value and temporal stability were validated using the polystyrene microsphere with a diameter of 2 μm (**Supplementary Figure S1**).

### Label-free live-cell imaging with TIE-based QPI (QPI-Mesoscope)

We first apply the quantitative phase imaging module of the Omni-Mesoscope for label-free live-cell imaging, where an incubator system was set up with the temperature set to 37°C, humid air, and 5% CO_2_. As shown in **Fig. 3A**, the quantitative phase images of cancer cells (SW480) exhibit significantly higher contrast compared to the conventional bright-field images, crucial for the identification of subcellular structures. Leveraging the mesoscale FOV, the QPI-Mesoscope can simultaneously monitor approximately 3000 cells with the ability to capture the heterogeneous cellular dynamics and rare events in a large cell population without the need for scanning or stitching various regions of interest. Moreover, for long-term monitoring, our snapshot imaging feature proves invaluable, as it substantially reduces light exposure, thereby minimizing the effects of phototoxicity (*17*). As shown in **Fig. 3B**, our system can track dynamic changes in the cytoskeleton with great sub-cellular details to visualize large skeletal structural features. We also observed rapid events of mitosis (**Fig. 3C**) that lasted for about 5 minutes over a 5-hour observation period.

**Figure 3.**
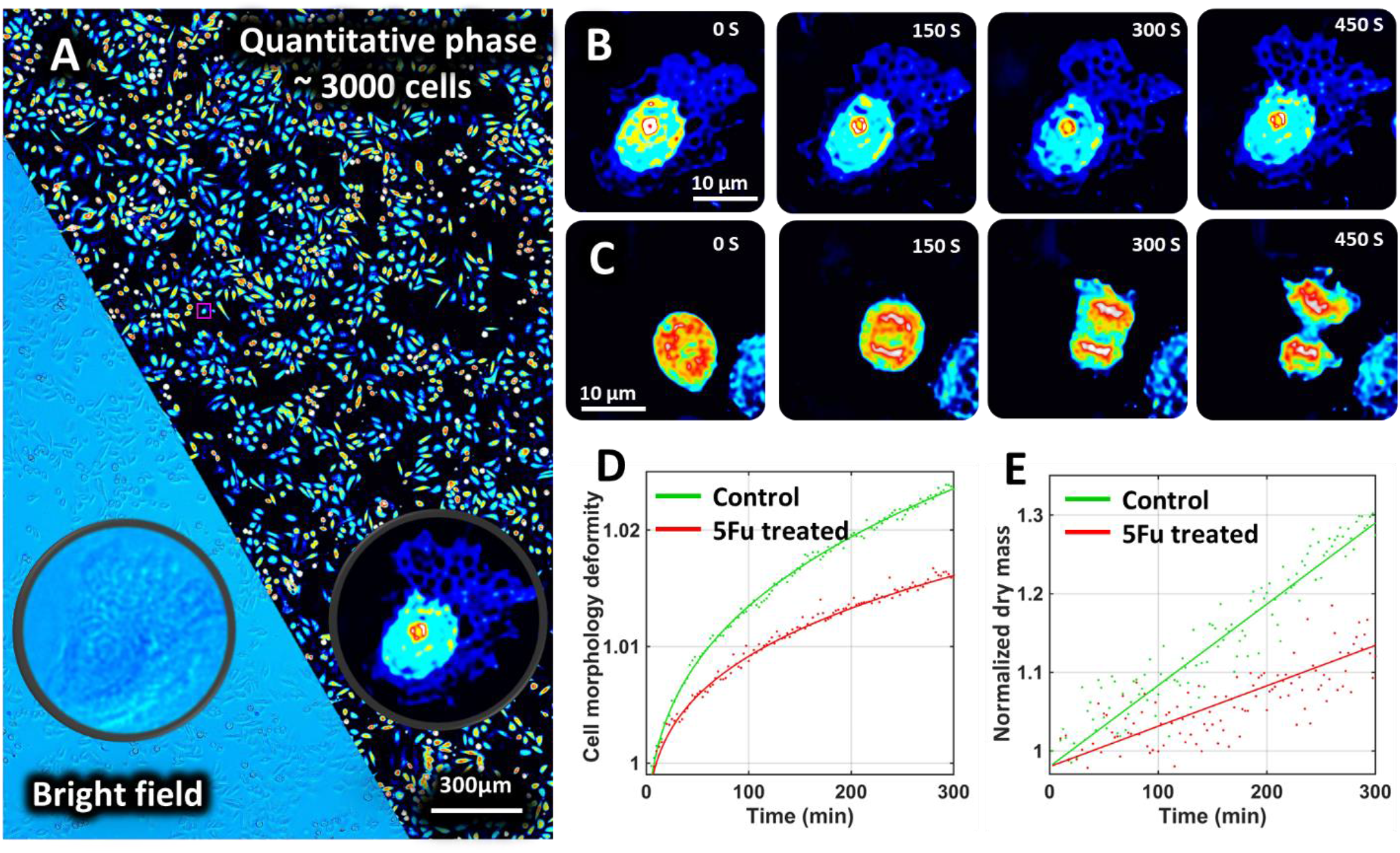
Label-free live-cell imaging with the quantitative phase imaging module of the Omni-Mesoscope. (A) Quantitative phase images of cancer cells in a single field of view, with approximately 3000 cells. The insert shows the comparison between the bright-field and quantitative phase images. (**B-C**) The dynamic changes of cytoskeleton and mitosis. (**D-E**) The quantification of cell morphological similarity and changes in dry mass in cancer cells without (control) and with the treatment of 5FU (5FU-treated) over 5 hours. Cell morphology deformity is quantified as the inverse of the image similarity between QPI images captured over 5 hours and those captured at the initial time point. The total dry mass of a cell is estimated by dividing the product of the quantifiable phase and cell area by the specific refractive index increment (*19*). Normalized dry mass is calculated by dividing the dry mass of the total cells captured at each time point by that captured at the initial time point.

This quantitative phase image provides enhanced image contrast and is also quantitative in nature (*16*). **Figures 3(D-E)** show that quantitative assessment of cell dry mass over the course of treatment with a chemotherapy drug, 5-Fluorouracil (5-FU). Interestingly, the results reveal that, as cells grow, their morphology gradually becomes less similar over time. In response to 5-FU treatment, the cell morphology undergoes less significant deformation. Similarly, the dry mass gradually increases during cell growth. The treatment with 5-FU also slows down the increase in dry mass, suggesting that the chemotherapy drug reduces protein synthesis during cell growth, consistent with the previous literature report (*18*).

### High-resolution multiplex fluorescence imaging of biological samples (Fluo-Mesoscope)

Next, we evaluated the imaging performance of Omni-Mesoscope for highly multiplex fluorescence imaging on biological samples using cancer cells (SW480). Three sub-cellular structural components—tubulins, lamin A/C, and centromeres—were utilized in this evaluation, as shown in **Fig. 4**. The results further illustrate the capability of Fluo-Mesoscope to obtain high-resolution fluorescence images over an ultra-large FOV at high throughput, while still resolving fine biological structures, such as the two closely adjacent centromeres. As shown in **Fig. 4(E)**, the Full Width at Half Maximum (FWHM) of a single centromere was measured at 0.7 μm and the distance between the two adjacent centromeres was at 1.5 μm, confirming the high-resolution imaging capability of Fluo-Mesoscope. These results also agree with the resolution defined using fluorescence microspheres (**Fig. 2**), further validating the accuracy and consistency of our mesoscopic imaging system.

**Figure 4.**
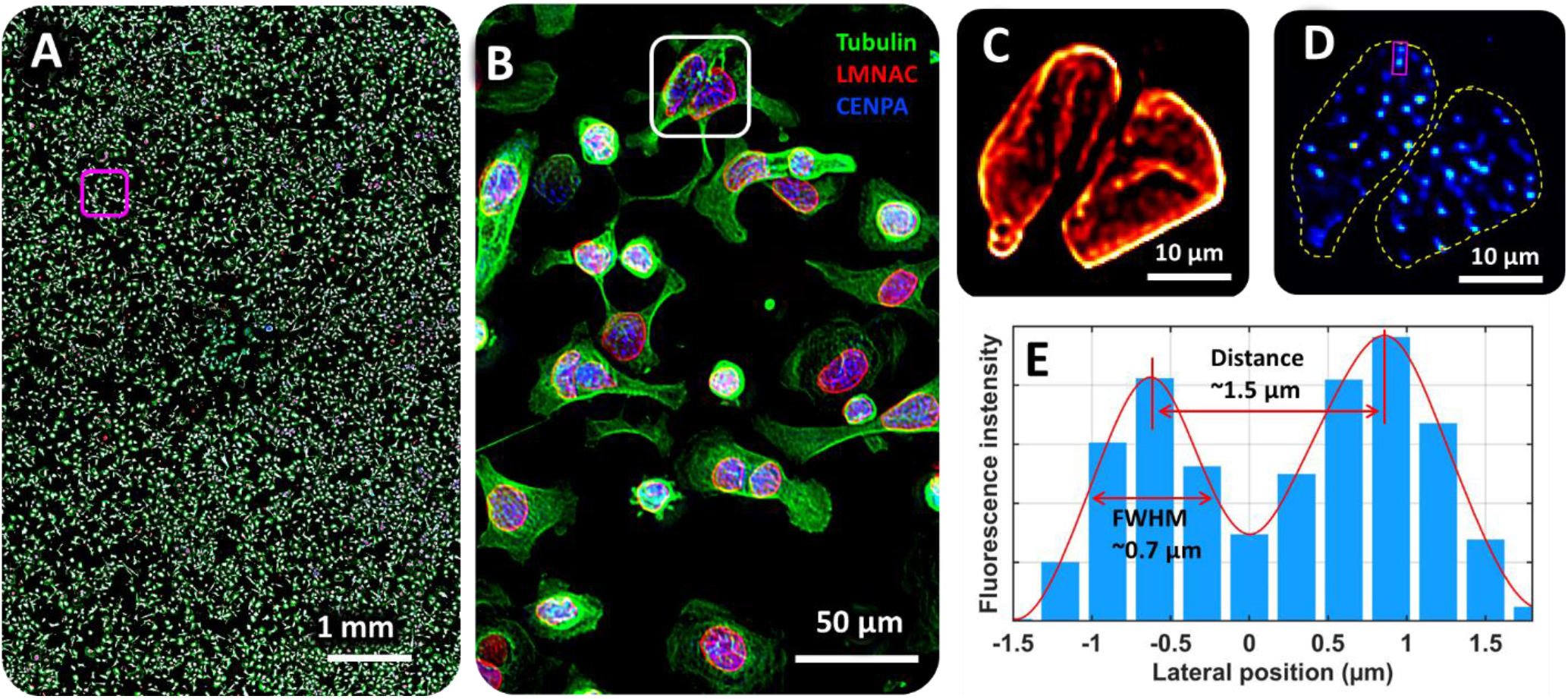
Performance of Omni-Mesoscope for multiplex fluorescence imaging of biological samples. (**A-B**) Fluorescence images of tubulin, lamin A/C, and centromeres over a large area (2×2 images stitched together), and (B) the zoom region of the magenta box from (**A**). (**C-D**) The zoom region of (**C**) lamin A/C (LMNAC) and (**D**) centromere from the white box in (**B**). (**E**) The lateral distribution of fluorescence intensity across the two adjacent centromeres (CENPA) in (**D)**, was measured at 1.5μm apart with a width of 0.7 μm.

### Omni-Mesoscope links the dynamic cell behaviors of rare events with underlying molecular characteristics

A key advantage of an ultra-large FOV in Omni-Mesoscope is its ability to detect rare dynamic cellular events, which can be linked to their underlying function through high-content molecular profiling *in situ*. This capability was demonstrated through the observation of the response of a cancer cell line, SW480, to the treatment of 5-fluorouracil (5-FU). Utilizing the capability of Omni-Mesoscope for quantitative phase imaging, dynamic morphological changes were simultaneously monitored for 5 hours across a large cell population of a few thousand cells within a single FOV. Within the predominantly slow-moving cell population, we identified a distinct subset that exhibited rapid and significant local movements. These observations are exemplified by a series of snapshot images in **Figs. 5(A1-A5)**, with representative videos (**Supplementary Videos 1-3**) providing further illustration. The temporal standard deviation map of phase images, presented in **Fig. 5(C)**, identifies a group of cells characterized by significant movement. As shown in **Fig. 5(B)**, the quantitative dry mass from the nuclei of these fast-moving cells appearred to be almost doubled compared to that of slow-moving cells. To uncover the function and molecular characteristics of these cells with distinct fast dynamics, we fixed the cells with paraformaldehyde (PFA) after five hours of live-cell imaging. As shown in the quantitative phase image (**Fig. 5(A6)**), some morphological changes are expected after fixation, such as cell shrinkage and flattening cytoplasm. However, the cellular morphology and the relative phase difference for different cells and sub-cellular compartments were largely preserved. Subsequent cyclic immunofluorescence staining and imaging on the Omni-Mesoscope revealed distinct molecular signatures from 14 different molecular markers for the same cells, presented in **Figs. 5(D1-D14)**. Notably, the fast-moving cells displayed significantly elevated levels of DNA damage marked by γH2AX, spread over the entire nucleus. This type of pan-nuclear pattern of γH2AX without foci indicates excess DNA damage from acute apoptosis (*20, 21*), distinct from the typical distinguishable γH2AX foci of DNA damage marker. Further, these cells showed increased levels of apoptosis (cleaved caspase-3), and proliferation (evidenced by Ki-67 and MCM7), alongside increased expression of cytoskeletal proteins (actin and tubulin), enhanced metabolic activity (ACC1, a marker for lipid or fatty acid metabolism), and higher DNA content (indicated in H3, HP1, H3K9ac, and centromeres), while similar levels of vimentin indicate the cells have not transitioned to a mesenchymal phenotype.

**Figure 5.**
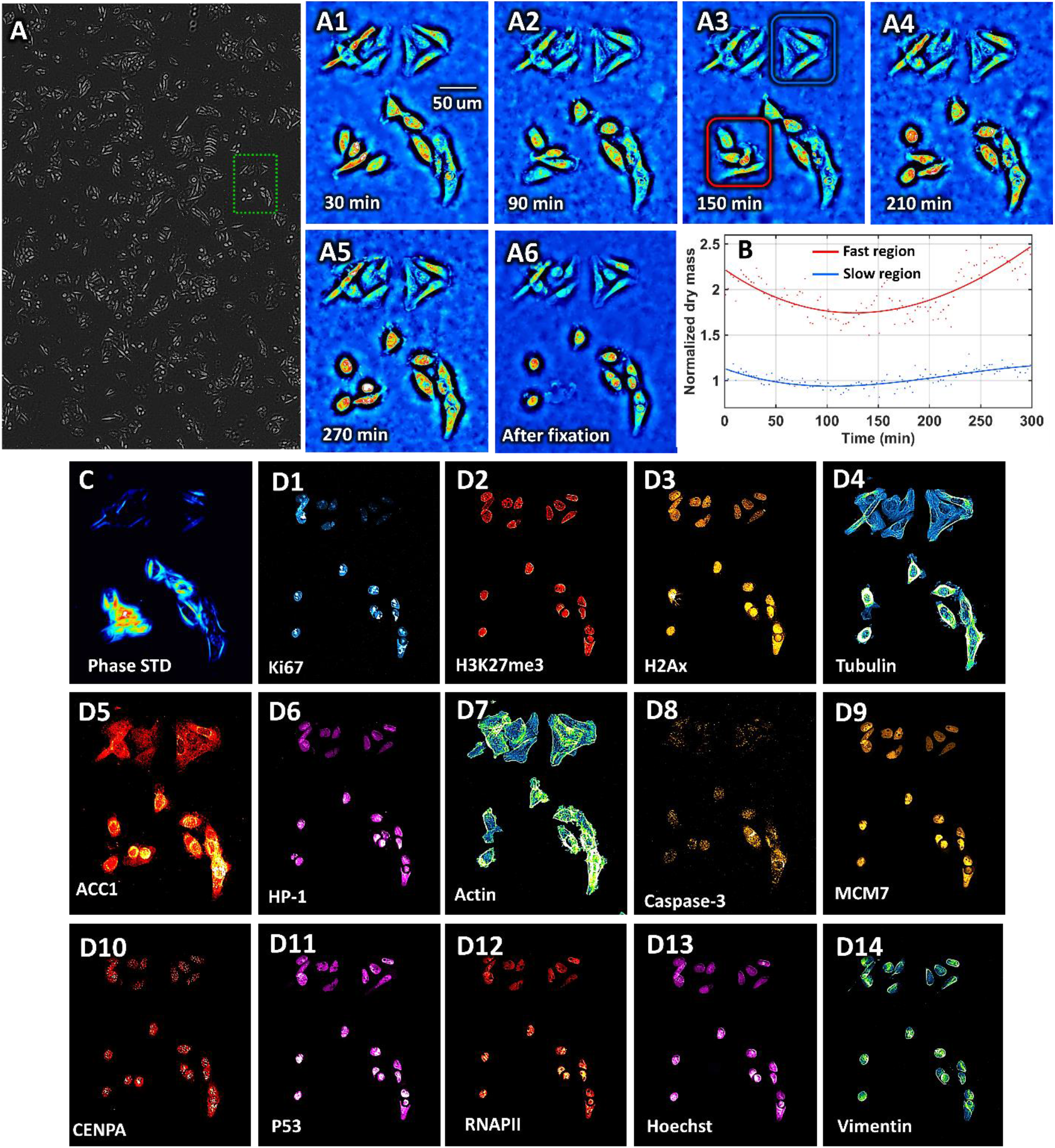
Correlation of dynamic cellular behaviors with molecular characteristics in SW480 cells treated with 5-FU. **(A)** The single FOV quantitative phase image of SW480 cells treated with 35μM 5FU for 3 days. (**A1-A5**) Representative sequential snapshot images from the green box in (**A**), which illustrate the dynamic morphological changes over 5 hours in SW480 cancer cells treated with 5FU for three days, captured within a single field of view encompassing thousands of cells. Most cells display minimal movement, while a subset exhibits significant local motion. (**A6**) Quantitative phase image post-fixation with paraformaldehyde (PFA), serving as a baseline for molecular analysis by preserving cellular structures for detailed examination. (**B**) The normalized dry mass of four cells in the fast-moving region and slow-moving region in (**A3**). (**C**) Temporal standard deviation map of phase images highlighting cells with pronounced movement, indicating areas of active cellular dynamics in response to treatment. (**D1-D14**) Fluorescence images of 14 molecular markers in the identified fast-moving cells and adjacent slow-moving cells. These markers include DNA damage (γH2AX), apoptosis (cleaved caspase-3), proliferation (Ki67, MCM7), cytoskeletal integrity (actin, tubulin), fatty acid metabolism (ACC1), tumor suppressor (p53), DNA content (H3), heterochromatin (HP1), euchromatin (H3K9ac), centromeres (CENPA), active transcription (phosphorated RNAPII) and mesenchymal marker (vimentin), providing a comprehensive molecular profile associated with the observed dynamic behaviors.

These observations collectively suggest that the fast-moving cells are undergoing significant DNA damage with features of pan-nuclear DNA damage, leading them toward lethal apoptosis in DNA replication stress. However, before succumbing to cell death, these cells show considerable chromatin condensation and dramatic mobility, supported by a marked increase in metabolic enzymes and cell proliferation proteins. The observed increase in DNA content and dry mass in cells suggests ongoing DNA replication and protein synthesis, yet without subsequent cell division. These results, revealed by the Omni-Mesoscope, provide functional insights into the complex cellular dynamics linked to their response mechanisms to chemotherapy drugs including those rare events.

Furthermore, the Omni-Mesoscope not only facilitates the detection of rare events but also enables a comprehensive analysis of the relationship between these events and the molecular characteristics of individual cells within the entire cell population, all obtained *in situ*. We conducted a Uniform Manifold Approximation and Projection (UMAP) analysis (*22*) for a comprehensive set of 20 markers on approximately 15,000 SW480 cells, both untreated and treated with 5-fluorouracil (5FU). The UMAP analysis in **Fig. 6** unveiled a distinct, small cluster of 5-FU-treated cells located on the right side of the visual representation. The UMAP revealed that there is a distinct cell cluster with exceptionally strong signals of γH2AX, indicative of severe pan-nuclear DNA damage. Notably, even within this cluster, there is significant heterogeneity in other morphological and molecular characteristics, such as nuclear size, cell dry mass, DNA content, proliferation status, transcription activities, and epigenetic states, suggesting diverse cellular outcomes in response to 5-FU treatment.

**Figure 6.**
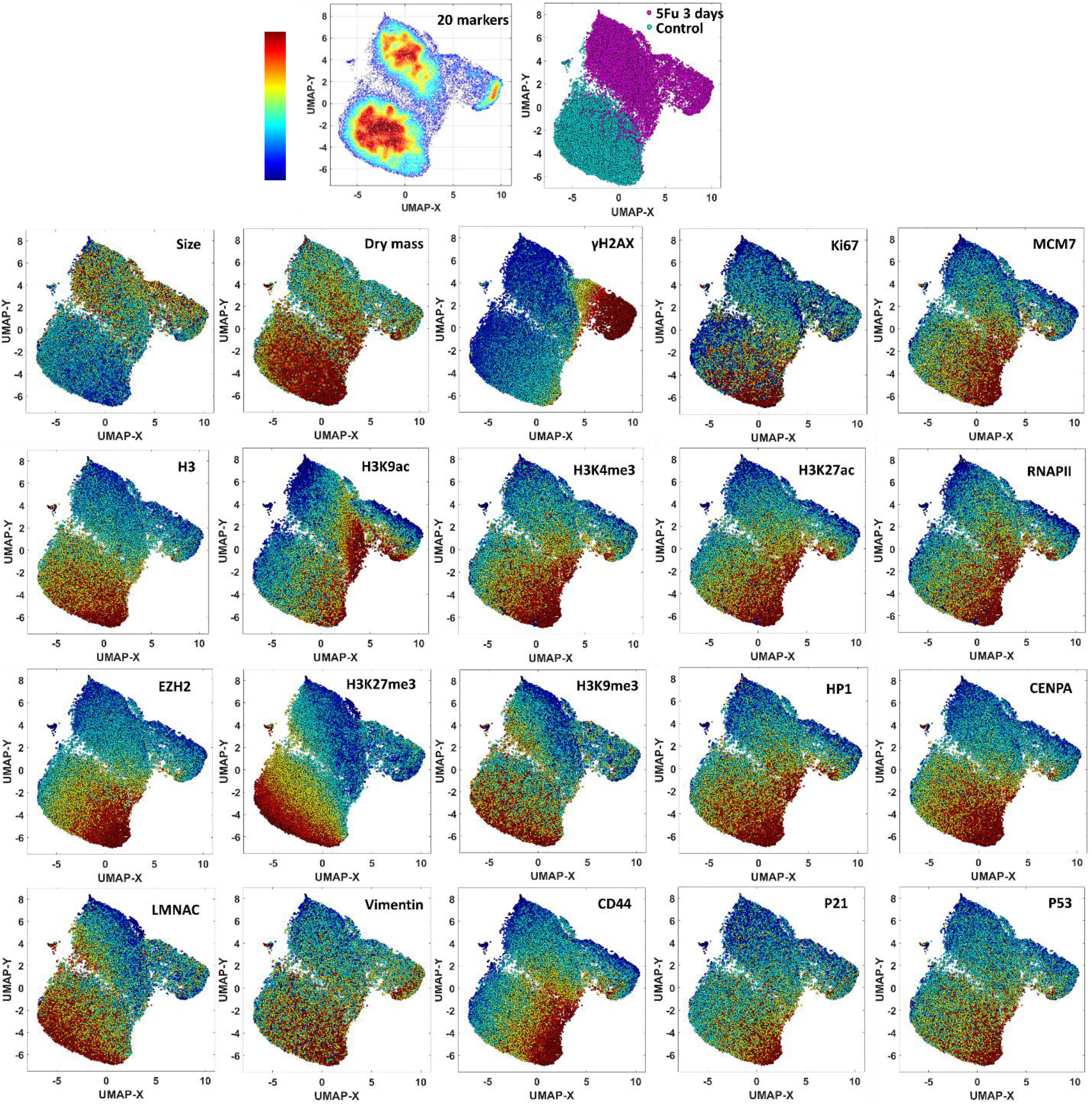
UMAP analysis for a total of morphological and molecular markers on ∼15,000 SW480 cells without (control) and with 5-FU treatment. Each panel represents the distribution of the fluorescence intensity for each marker on the clustered UMAP. The MATLAB UMAP package is used for data analysis with default parameter setting (min_dist = 0.3, n_neighbors = 15, metric = Euclidean, randomize = 1).

Further analysis also revealed a strong correlation between the temporal standard deviation map of phase images from the fast-moving cells and those displaying intense γH2AX signals, as shown in **Supplementary Figure S2**. Interestingly, those fast-moving cells represent a minor fraction (<10%) of the cell population, even among those with excessive DNA damage, emphasizing the rarity of these events.

### Omni-Mesoscope reveals the formation of polyploidy in drug-resistant cancer cells and their underlying molecular characteristics

The next example highlights the use of the Omni-Mesoscope in characterizing drug-resistant cancer cells. We conducted live-cell imaging of drug-resistant cancer cells that adaptively developed resistance throughout the treatment with increasing concentration of 5-FU over approximately 3 months. During a 5-hour imaging session, we monitored the cellular dynamics in these drug-resistant SW480 cells in the presence of 6μM 5-FU and compared them to the dynamics of the corresponding parent SW480 cancer cells. As the cancer cells adapt and develop chemoresistance, the large FOV provided by Omni-Mesoscope allowed us to observe the highly heterogeneous dynamic behaviors of these cells. In particular, we observed the formation of polyploid cells through the mechanism of cell engulfing and cell fusion. As shown in **Figs. 7(A1-A12)**, a cell was first engulfed into an existing polyploid cell, becoming flattened—a state corroborated by a lower phase value indicative of a thinner cell inside the polyploid cells. This engulfed cell was then divided into two daughter cells within the large polyploid cell (**Supplementary Video 4**). Three such cell-engulfing events (**Supplementary Videos 4-6**) were observed over the entire FOV of ∼5 mm^2^ with approximately 1500 cells over 5 hours. Additionally, an event involving cell fusion with the polyploid cell was observed (**Supplementary Video 6**).

**Figure 7.**
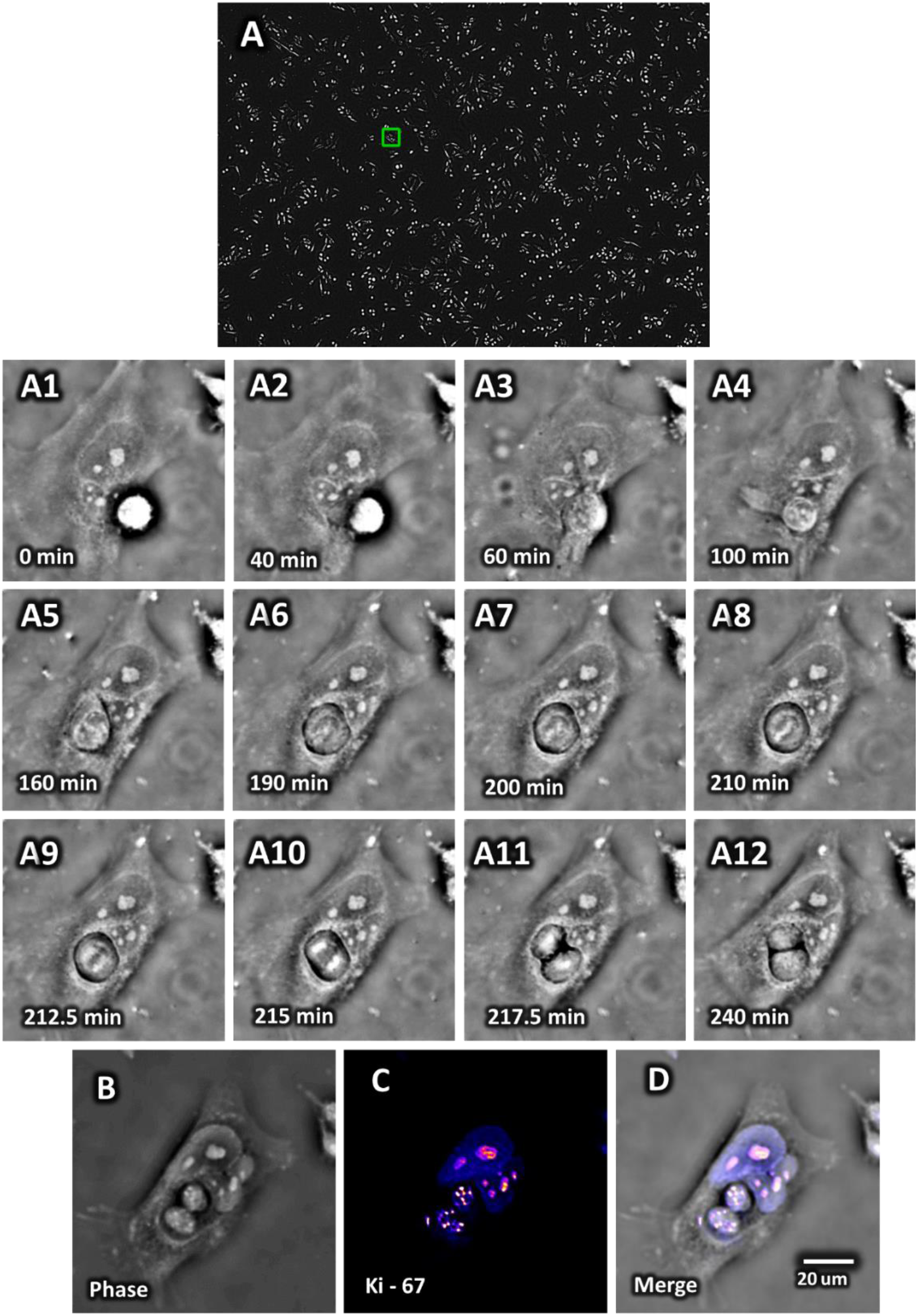
Formation of polyploid cells in drug-resistant cancer cells. **(A)** The single FOV quantitative phase image of drug-resistant cancer cells SW480 in the presence of 6μM of 5-FU. **(A1-A12)** Representative sequential images from the green box in (A), reveal the formation of polyploidy in drug-resistant cancer cells. **(B)** Quantitative phase image after fixation, serving as a baseline for molecular analysis by preserving cellular structures for detailed examination. (**C**) Fluorescence images of Ki-67 markers. (**D**) The merged quantitative phase and fluorescence image show a high level of correlation between the phase value and the fluorescence intensity of Ki-67.

Importantly, the sub-micron resolution of Omni-Mesoscope enables the detection of functionally significant sub-cellular dynamics. As illustrated in **Figs. 7(B-D)**, the quantitative phase image reveals dense puncta within the cell nuclei, which correlate strongly with the corresponding Ki67 fluorescence intensity. This observation indicates the label-free quantitative phase images can reveal the dynamic morphological changes with sub-cellular details, specifically within nuclear protein Ki67. Smaller dense puncta of Ki67 are evident in the daughter cells derived from the engulfed cell, while larger dense Ki67 puncta are observed in the nuclei of the existing polyploid cells, reflecting their different stages of the cell cycles (*23*).

Building on these observations, **Fig. 8** details the underlying high-content molecular profiling that outlines the specific molecular characteristics of these polyploid cells. These cells exhibit strong stemness (high level of CD44), active fatty acid metabolism (ACC1), significant proliferation (MCM7), and high level of DNA damage (γH2AX). These molecular traits are consistent with the proposed mechanism whereby polyploidization enables cancer cells to survive harmful conditions (*24*). The previous studies also reported cell cannibalism and cell fusion as possible mechanisms for polyploidy formation (*24*), which our direct observations using dynamic cellular imaging in a single FOV have confirmed. Further, the quantitative phase images captured highly dynamic cytoskeleton activities, such as cytoskeleton extrusion, which is particularly pronounced in these polyploid cells. The underlying molecular profiling confirms that the extrusion structure comes from both actin and tubulin, suggesting their role in facilitating fast cell movement.

**Figure 8.**
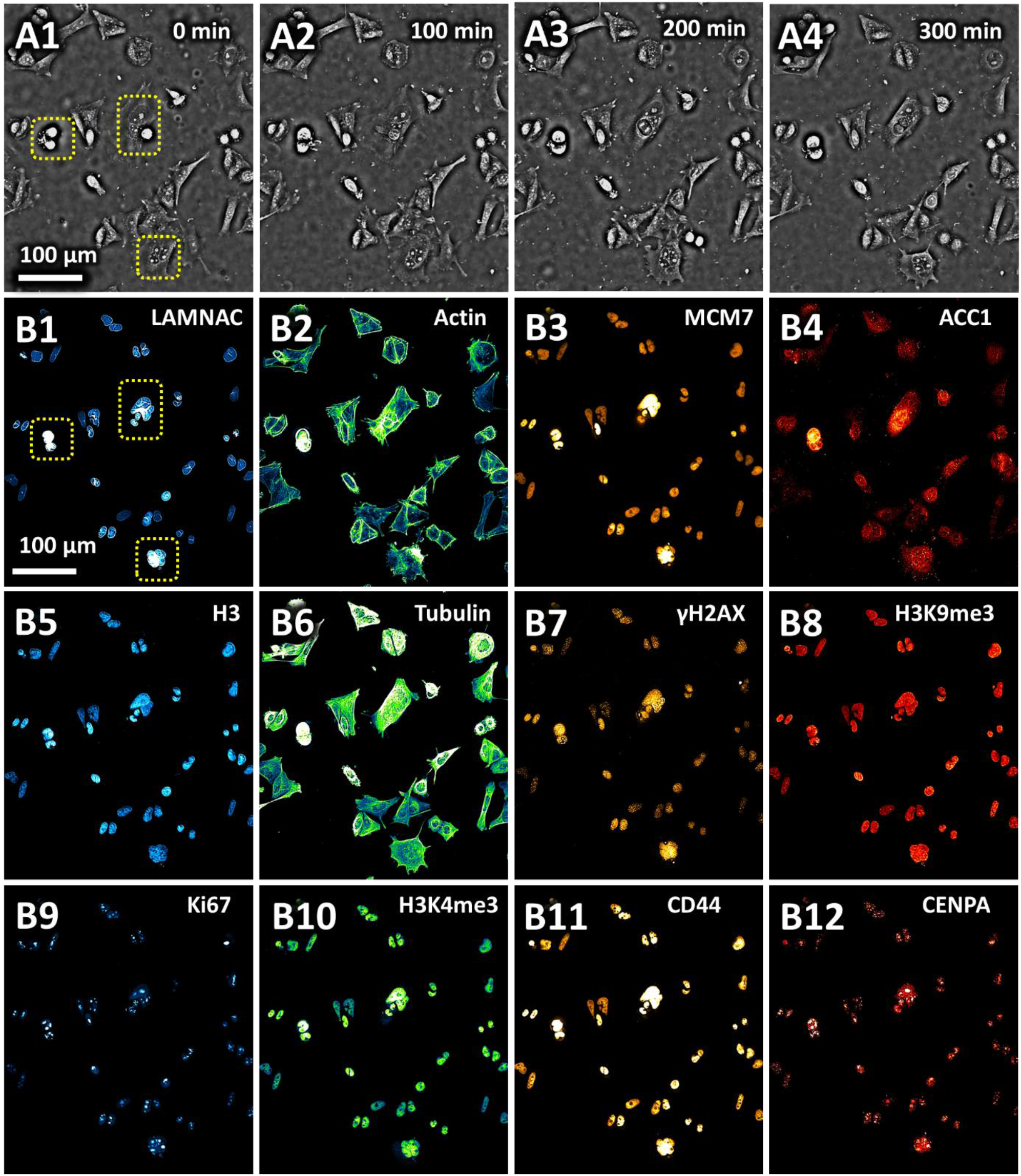
High-content molecular profiling of polyploid cells in the drug-resistant SW480 cells. (**A1-A4**) Representative sequential images cropped from the FOV, illustrate the dynamic morphological formation of polyploidy over 5 hours in SW480 drug-resistant cells. The yellow dashed box highlights the polyploid cells. (**B1-B12**) Fluorescence images of 12 molecular markers in the polyploidy cells and other drug-resistant SW480 cells. These markers include DNA damage (γH2AX), proliferation (Ki67, MCM7), cytoskeletal integrity (actin, tubulin), fatty acid metabolism (ACC1), DNA content (H3), heterochromatin (H3K9me3, H3K4me3), centromeres (CENPA), lamin A/C (LMNAC), cancer stemness (CD44), providing a comprehensive molecular profile associated with the observed polyploidy.

This approach underscores the value of combining label-free dynamic cellular imaging with deep molecular analysis to uncover critical cellular responses to treatment. By facilitating the identification of specific cell subsets undergoing significant stress or damage, such as those highlighted in the UMAP analysis, our methodology that coupled morph-dynamic imaging over a large cell population with subsequent high-content molecular profiling, offers a powerful tool for understanding the heterogeneity of cellular responses to pharmacological interventions.

### High-throughput 3D volumetric super-resolution imaging of thick tissue (Ex-mesoscope)

As a versatile imaging platform, our Omni-Mesoscope can also enhance the throughput for highly multiplexed tissue imaging. As shown in **Supplementary Fig. S4**, with significantly improved FOV and imaging speed, the whole-slide imaging of a tissue section from a small tissue biopsy only requires two images in just a few seconds.

A notable limitation in this ultra-large field of view (FOV) wide-field optical microscopy lies in its limited 3D sectioning ability, evidenced by an axial resolution of approximately 6 μm (**Fig. 2**) in Omni-Mesoscope. The long depth of field increases the background from thick tissue, which significantly degrades their image quality. To enable 3D volumetric imaging, we have integrated Magnify (*25*), the state-of-the-art variant of expansion microscopy into our workflow. Expansion microscopy physically enlarges biological specimens, enabling imaging at resolutions surpassing those attainable with conventional microscopy methods. Crucially, the expansion process not only enhances resolution but also significantly mitigates scattering and increases sample transparency.

As shown in **Fig. 9(A)**, the 3D volumetric super-resolution images of the expanded samples can be obtained with Omni-Mesoscope (estimated to be ∼150 nm lateral resolution and 1.5 μm axial resolution based on 4.5 expansion ratio), on a 30μm-thick section of mouse small intestine tissue. A comparative analysis of tissues, both pre-expansion (**Figs. 9(B, D)**) and post-expansion (**Figs. 9(C, E)**), reveals a marked enhancement in image resolution post-expansion. This enhancement allows for the clear visualization of condensed chromatin foci within nuclei and enables the precise axial localization of individual cells.

**Figure 9.**
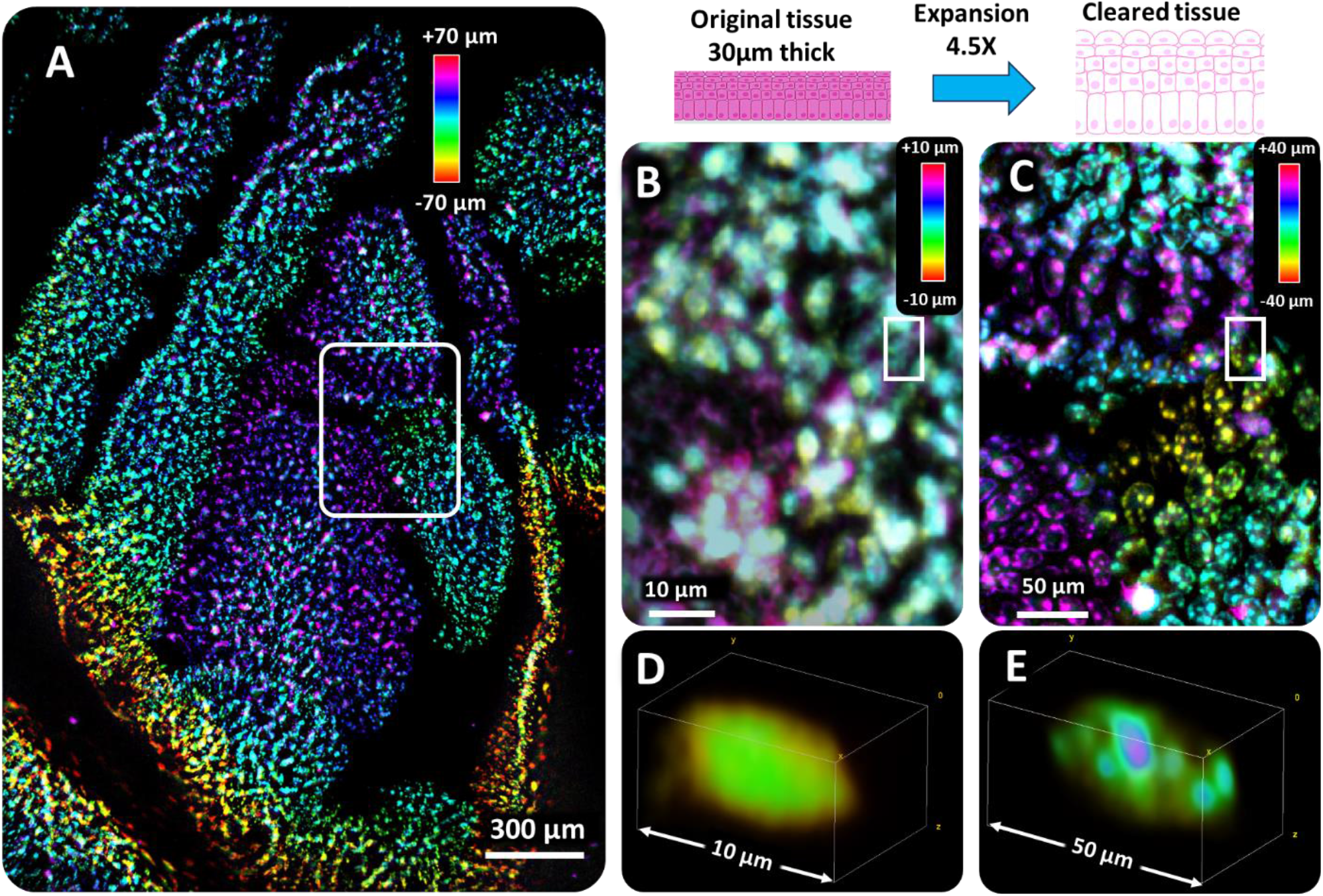
Three-dimensional (3D) Volumetric Super-Resolution Imaging of Thick Tissue Using the Omni-Mesoscope with Expansion Microscopy. (**A**) 3D volumetric super-resolution image of nuclei (DAPI) on an expanded sample (∼4.5x expansion, 1XPBS) from a 30μm-thick section of mouse small intestine tissue, obtained using the Omni-Mesoscope. (**B, D**) Images of the tissue section before expansion, showing the degraded resolution and high background. (**C, E**) Images of the tissue section post-expansion, demonstrating significant improvements in image resolution and 3D sectioning capability of Omni-Mesoscope.

It is important to note that the Omni-Mesoscope offers a significant increase in imaging throughput for expansion microscopy. This advantage is particularly important given that expanded samples come with a dramatic volume increase—up to three orders of magnitude post-expansion. Consequently, the synergy between the Omni-Mesoscope and expansion microscopy shows tremendous potential for ultra-high-throughput 3D volumetric super-resolution imaging of thick tissue samples, to explore complex biological structures with unprecedented clarity and detail at high speed.

## DISCUSSION

We present Omni-Mesoscope, a multiscale multi-modal imaging system to address the limitations of traditional wide-field optical microscopy. Omni-Mesoscope not only provides high-throughput, high-resolution imaging, but also utilizes label-free quantitative phase imaging to observe cell dynamics across a large cell population. Moreover, it uniquely connects the non-invasive monitoring of morphological dynamics with information-rich molecular characteristics at an unparalleled scale, resolution, and functional insights. Technically, the Omni-Mesoscope achieves a sub-micron lateral resolution for the entire field of view of ∼5mm^2^, while using off-of-shelf, cost-effective objective lenses and a large-format camera, at a cost of around $4000, making it highly scalable and affordable. Additionally, it is compatible with the standard coverslip-based live-cell imaging chamber without the need for a specialized immersion-based imaging chamber.

We demonstrate the transformative potential of the Omni-Mesoscope in characterizing cellular responses to chemotherapy and chemoresistance. By capturing specific dynamic events over a large cell population, this mesoscale quantitative phase imaging system reveals dynamic sub-cellular changes along with their underlying molecular characteristics, marking a significant advancement in in cell biology research. Detailed live-cell imaging and molecular profiling provide a comprehensive understanding of cellular responses to 5-fluorouracil treatment. The identification of a subset of SW480 cells exhibiting rapid movements and significant DNA damage highlights the potential of Omni-Mesoscope in providing the functional interpretation of specific dynamic cell behaviors. The dramatically increased cell mobility could be one enabling capability for those cells to eventually detach from the substrate and undergo apoptosis.

Additionally, Omni-Mesoscope facilitates direct observation of cell cannibalism and cell fusion in the formation of polyploid cells in drug-resistant cancer cells, all within a single FOV. Such dynamic events are then correlated with the molecular traits characteristic of highly aggressive cancer cells such as increased stemness, proliferation, and mobility. Consequently, the integration of mesoscale quantitative phase imaging of cell dynamics together with high-content molecular profiling at sub-micron resolution significantly enhances our ability to dissect the heterogeneity of cellular responses to therapeutic agents and opens new possibilities to probe the complex interplay between cellular dynamics and molecular mechanisms in real-time over a large cell population.

The introduction of expansion microscopy within our imaging workflow further enables 3D volumetric super-resolution imaging of thick tissue samples. The Omni-Mesoscope, when coupled with expansion microscopy, shows significantly improved image resolution and 3D sectioning capability on thick tissue sections. It also addresses a major bottleneck in imaging throughput for 3D imaging of cleared and expanded samples. Future work to further explore the synergy between the Omni-Mesoscope and expansion microscopy will enable ultra-high throughput 3D volumetric super-resolution imaging of thick tissue samples to explore complex tissue architecture and molecular characteristics with unprecedented clarity and detail at high throughput.

In conclusion, the Omni-Mesoscope, with its integration of quantitative phase microscopy and highly multiplex fluorescence imaging, presents a significant advancement in our ability to simultaneously capture and analyze the dynamic and molecular essence of cells in unprecedented detail and high molecular content. In the context of the existing mesoscope and light-sheet imaging systems, most target fluorescence imaging with a lateral resolution of 2-5 μm (*4*–*10*).

While the customized mesolens have a similar NA as our system, scaling it up has been a major challenge due to the high cost and labor-intensive nature of manufacturing such a mesolens (*26*). Compared to Fourier Ptychographic Microscopy (FPM) (*27*), our Omni-Mesoscope achieves a shorter depth of focus (6-8 μm) and a higher temporal acquisition speed of 4 fps, offering one-order-of-magnitude improvement in optical sectioning and temporal resolution. It also integrates seamlessly with fluorescence imaging. Our Omni-Mesoscope fills in a gap in the field by simultaneously capturing large-scale temporal dynamics and sub-cellular details over a large cell population while also delivering the high-content molecular characteristics required to decode heterogeneous responses among individual cells in distinct states. We recognize that the Omni-Mesocope, in its current form, has limited performance in imaging 3D scattering samples, due to limited axial resolution. Future efforts will address this limitation via parallel scanning, computational imaging, and artificial intelligence-assisted image enhancement. This system opens new avenues for the exploration of biological systems, potentially revolutionizing our approach to diagnostic and therapeutic strategies in the biomedical sciences.

## MATERIALS AND METHODS

### Instrument Development of Omni-Mesoscope

We built the Omni-Mesoscope using cost-efficient off-the-shelf components. In brief, we adopted six cost-efficient industry-grade lasers from Civil Laser to provide a large FOV illumination (*28*): 405nm (LSR405CPD-1.2W), 470nm (LSR470SD-2.5W), 530nm (LSR530SD-1.2W), 589nm (LSR589H-1W), 650nm (LSR650SD-1W), and 750nm (DL750-T2-1.5W).(*29*) These six laser lines were combined with dichroic mirrors (T420lpxr, T495lpxr, T556lpxr, T610lpxr, T685lpxr, Chroma), coupled into a multimode fiber (M97L02, Thorlabs) for beam shaping into a flat illumination field (*29, 30*). The square core multimode fiber transforms the original beam with an irregular shape into a uniform beam with a square shape. A high-frequency vibration motor (VZ6DL2B0055211, Digikey) was used to reduce the laser speckle and achieve a high uniformity of up to 99%. The laser intensity is electronically controlled by an Arduino board (Arduino Nano Every) with TTL mode.

To increase the signal-to-noise ratio (SNR), we decoupled the illumination path and detection path, and the light was obliquely introduced from the bottom of the sample with an incident angle of 60°, which is larger than the collection angle of the objective lens (45°). This configuration minimizes scattered and reflected light entering the detector, thus significantly reducing the background noise. A custom motorized filter wheel with 9 band-pass emission filters (#67-040-438CWL, #87-762-510CWL, #87-765-549CWL, #84-116-572CWL, #87-767-615CWL, #33-328-631CWL, #86-996-676CWL, #67-052-692CWL, #84-123-832CWL) was used to switch between different fluorescence imaging channels.

Fr optimal live-cell imaging, we designed a tabletop incubator with two levels of environmental control to ensure a stable environment. We used a precise inner chamber incubator (TCS-200, Amscope) to maintain the cell chamber at 37°C, and the humid air with 5% CO_2_ was injected directly into the chamber incubator. The outer incubator includes a temperature controller (ITC-308, Inkbird) and a high-power heater (WY-H1-B1, YOUCIDI) to maintain the temperature of the whole tabletop space consistently at 37°C with minimal fluctuation (<0.2°C).

### Image acquisition procedures

Our Omni-Mesoscope includes two distinct imaging modes: QPI-Mesoscope mode for label-free live-cell imaging and Fluo-Mesoscope mode for multiplex fluorescence imaging. For live-cell imaging mode, we first turn on the incubation system for appropriate temperature (37°C) and CO_2_ concentration (5%). Our incubation system achieves a stable condition in about 10 minutes. We then mount the imaging chamber and start the automatic live-cell imaging. During the imaging process, the Mesoscope automatically finds the focus by scanning the sample along the axial direction and identifying the focal plane where the captured bright-field images exhibit the lowest normalized variance of the pixels (*31*). The bright-field images captured at different axial positions at ±2 μm step size are also used for TIE-based quantitative phase reconstruction and refocusing. Live cells were monitored for over 5 hours with a time interval of 150 seconds. For multiplex fluorescence imaging mode, a similar bright-field image-based strategy was used for autofocusing. Furthermore, we also captured a 3D stack of fluorescence images for subsequent region-based refocusing. The exposure time was adjusted to achieve a high contrast for each fluorescence channel.

### Image processing and statistical data analysis

In this study, three major image processing steps (Supplementary Figure S3) were used to achieve state-of-the-art image quality.

#### Flat field calibration

Two strategies were used to achieve uniform fluorescence imaging. First, a multimode fiber with a square core was utilized to create uniform illumination, with a vibration motor employed to reduce speckle effects (*29*). This approach enables illumination uniformity of over 96%. However, the detection efficiency of the objective lens was not uniformly distributed. Second, under uniform illumination, multiple fluorescence images at different positions of the same sample were captured. The average fluorescence intensity at each position was calculated to generate a 2D relative intensity map. Subsequently, a 2D cubic polynomial function was fitted to this map to obtain a 2D normalization map for each pixel. This map calibrated the detection efficiency and produced the final optimized flat field image with uniformity exceeding 96%.

#### Region-wise refocusing

Given that all objective lenses exhibit field curvature, region-wise refocusing was implemented to address defocusing issues across a large field of view.

Fluorescence microspheres placed on a #1.5 coverslip were used to capture 3D image stacks, from which the 3D position of each emitter was retrieved. A 2D cubic polynomial function was then fitted to generate a field curvature map. The full image was divided into 9×6 regions, and refocusing was performed by identifying the focal plane within each region. In-focus regions replaced defocused ones to minimize defocus distortions, resulting in a reduction of defocusing distance to ±1 μm.

#### Background reduction, denoising, and deconvolution

The rolling ball algorithm (implemented with the MATLAB function ‘imboxfilt’) with a 20-pixel diameter was utilized to reduce background, leading to an average signal-to-background ratio improvement of 2-5 folds (*32, 33*). Additionally, a Wiener filtering algorithm (implemented with the MATLAB function ‘wiener2’) was applied to reduce noise and enhance spatial resolution, resulting in a 10-20% improvement (*32, 34*).

#### UMAP analysis

The UMAP package written in MATLAB (*35*) is used for data analysis with default parameter setting (min_dist = 0.3, n_neighbors = 15, metric = Euclidean, randomize = 1).

### Transport of intensity equation (TIE) based quantitative phase imaging (QPI-Mesoscope)

The Transport of Intensity Equation (TIE) is essentially an expression of the conservation of energy in optics (*12, 36*). It establishes a direct mathematical relationship between the spatial phase and the derivative of intensity along the optical axis, enabling the retrieval of quantitative phase values by measuring the phase-induced intensity gradient.

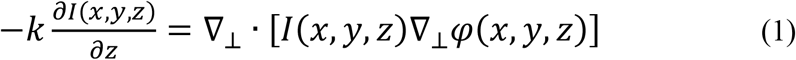

where ϕ is the phase, *k* is the wave number as 2π/*λ, λ* is the wavelength, *I* is the image intensity, 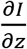 represents the intensity derivative, ∇⊥ is the lateral gradient operator, *x* and *y* indicate the lateral spatial coordinate, and *z* indicates the axial axis. For a phase object under uniform illumination, the intensity along the optical axis is nearly constant, the TIE can be further simplified as a Poisson equation in the following form:

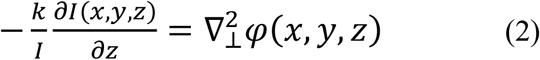

Then, the spatial phase distribution can be expressed as:

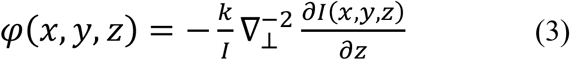

The inverse Laplacian operator (∇^−2^) can be efficiently calculated by using the Fast Fourier Transform (FFT). Therefore, the phase distribution can be reconstructed as:

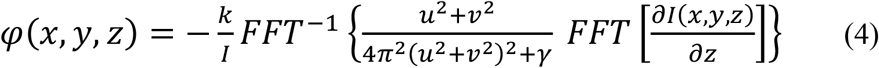

where *u* and *v* are the spatial frequencies corresponding to the lateral axis, and *γ* is a regularization parameter to avoid the singularity and remove noise-induced artifacts. Since the axial intensity derivative cannot be measured directly, finite differences are used to approximate the derivative in practice. In its simplest form, TIE phase retrieval requires only two bright-field intensity images collected at different focuses along the axis dimension.

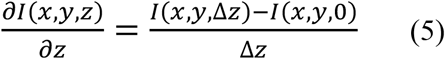

where Δz is a small defocus axial distance to provide a good estimation of the intensity derivative.

### Dry mass calculation

The total dry mass of a cell is estimated by dividing the product of the phase value and cell area by the specific refractive index increment (*19*). Normalized dry mass is calculated by dividing the dry mass of the total cells captured at each time point by that captured at the initial time point.

### Treatment of SW480 cells with 5-fluorouracil (5-FU)

The SW480 cells were exposed to 35μM 5-FU for durations of 3 days. Subsequently, the cells were imaged using QPI-Mesoscope under live-cell imaging conditions (37°C, 5% CO_2_) for 5 hours. Immediately after QPI, the cells were fixed with 4% paraformaldehyde (PFA) for 15 minutes, followed by cyclic fluorescence staining.

### Development of drug-resistant SW480 cells

The SW480 cells were gradually exposed to increasing concentrations of 5-FU, starting with 1 μM, followed by 2 μM, and then 4 μM, before reaching a final concentration of 6 μM. The cells were maintained at each concentration for about 4 weeks. Throughout this period, the culture medium at each concentration was refreshed every three days to maintain optimal conditions. Subsequently, the cells were imaged live using QPI-Mesoscope under 37°C and 5% CO_2_ for 5 hours. Immediately after imaging, the cells were fixed with 4% PFA for 15 minutes, followed by cyclic fluorescence staining.

### Cyclic immunofluorescence (CycIF) staining

Cultured cells were briefly washed with 1xPBS before being fixed with 4% PFA for 15 minutes at room temperature (RT). Fixed cells were rinsed with 1xPBS twice before permeabilization with 0.2% Triton X100 in 1xPBS for 10 minutes. The cells were incubated with a blocking buffer containing 5% BSA in 1xPBS for 30 minutes at RT. All antibodies were diluted in a blocking buffer. The dilutions and usages of antibodies in each cycle were documented (**Supplementary Table S1)**. The cells from the initial staining or previous bleached cycle were stained with diluted primary or fluorophore-conjugated antibodies at 4°C for 12 hours in a moisture chamber. The cells were then washed with 1xPBS three times.

For indirect immunofluorescence staining, after incubation with primary antibodies, diluted secondary antibodies in blocking buffer were added to the cells for incubation at room temperature for 2 hours, followed by three washes with 1xPBS. The stained cells were stored in 1xPBS until imaging.

### Fluorophore inactivation

After each round of fluorescence imaging, the fluorophores were chemically inactivated by incubation in a bleaching buffer containing 4.5% (w/v) H_2_O_2_ (Sigma, Cat. H1009), 20mM NaOH (solution made from pellets, Sigma-Adrich, Cat. S5881) in 1xPBS for 20 minutes. The samples were continuously illuminated by two high-power full-spectrum LEDs (YUJILEDS-CRI-95, 380nm-1000nm, 3.6W) for fluorescence photobleaching. After 1 hour of photobleaching, the cells were washed with 1xPBS three times to remove residual oxidation solution before being subjected to a subsequent round of immunofluorescence staining.

### Reagents and Methods for Magnify

Reagents used in this study included Paraformaldehyde (PFA, Sigma Aldrich, P6148), Ethanol (Pharmco, 111000200), Xylene (Sigma Aldrich, 214736), Sodium Acrylate (SA, AK Scientific, R624; Santa Cruz Biotechnology, sc-236893B), N-dimethylacrylamide (DMAA, Sigma Aldrich, 274135), Acrylamide (AA, Sigma Aldrich, A8887), N,N’-Methylenebisacrylamide (BIS, Sigma Aldrich, M7279), Tetramethylethylenediamine (TEMED, Sigma Aldrich, T9281), 4-Hydroxy-2,2,6,6-tetramethylpiperidine 1-oxyl (Sigma Aldrich, 176141), Sodium Chloride (NaCl, Sigma Aldrich, S6191), Phosphate Buffered Saline 10x solution (Fischer Scientific, BP399-1), Ammonium Persulfate (APS, Sigma Aldrich, A3678), Potassium Persulfate (KPS, Sigma Aldrich, 216224), Methacrolein (Sigma Aldrich, 133035), Ethylenediaminetetraacetic acid (EDTA, VWR, BDH7830-1), TritonX-100 (Sigma Aldrich, T8787), Tris-BASE (Fischer Scientific, BP152-1), Proteinase K (Fischer Scientific, EO0491), Sodium Dodecyl Sulfate (SDS, Sigma Aldrich, L3771), Urea (Sigma Aldrich, U5378), Glycine (Sigma Aldrich, G8898).

### SA Stock Solution Preparation

A 50% final concentration was achieved by adding deionized water in multiple increments, stirring continuously to ensure complete dissolution. Adequate time was allowed for complete dissolution before final volume adjustment.

### Monomer Solution Preparation

Comprised of 4% v/v DMAA, 34% v/v SA, 10% v/v AA, 0.02% v/v BIS, and 1% w/v NaCl in 1x PBS, the solution was stored at 4°C until use.

### Heat Denaturation Buffer Preparation

This buffer was made of 1% w/v SDS, 0.75% w/v Glycine, 8M Urea, 25 mM EDTA, and 500 mM Tris-BASE in 2x PBS, adjusted to pH 8.5 and stored at room temperature.

### Deparaffinization of FFPE Tissue

Formalin-fixed paraffin-embedded (FFPE) pathogen-infected tissue samples underwent deparaffinization through sequential immersion in 2x xylene, 2x 100% ethanol, followed by 95%, 70%, and 50% ethanol dilutions, and finally, doubly deionized water. Each step was carried out at room temperature, lasting 3 minutes.

### In Situ Polymer Synthesis

Prior to gelation, 0.2-0.25% w/v APS, 0-0.25% v/v TEMED, 0.001% w/v 4HT, and 0.1-0.25% v/v methacrolein were added to the monomer solution. After vortexing, the samples were incubated with the gelation solution for 5 to 40 minutes at 4°C to allow diffusion without premature gelation. Gelation was completed overnight in a humidified chamber at 37°C, using a setup made from spacers cut from #1.5 cover glass and a glass slide.

### Sample Homogenization and Expansion with Magnify

After gelation, mouse colon samples were immersed in a denaturant-rich buffer (1% w/v SDS, 8M Urea, 25 mM EDTA, 2x PBS, pH 7.5) at room temperature for 30 minutes. The homogenization was conducted in a pressure cooker at 120 °C for 80 minutes. After homogenization, samples were washed twice with 1% decaethylene glycol monododecyl ether (C12E10) in 1x PBS at 60°C for at least 15 minutes per wash, followed by two washes at 37°C to remove SDS.

### Post-Expansion fluorescence staining

Expanded samples were incubated with DAPI (1 μg/ml) for 10 minutes before imaging.

## Author contributions

Conceptualization: HM, YL

Methodology: HM

Investigation: HM

Formal analysis: HM, MC

Resources: JX, MC

Visualization: HM, MC, YL

Project administration: YL

Writing—original draft: HM

Writing—review & editing: HM, YL

## Data and materials availability

All data needed to evaluate the conclusions are shown in the paper, Supplementary Figures, Supplementary Videos and Supplementary Table.

## References

1. Y. Sasai, Cytosystems dynamics in self-organization of tissue architecture. Nature 493, 318–326 (2013).

2. A. Rao, D. Barkley, G. S. França, I. Yanai, Exploring tissue architecture using spatial transcriptomics. Nature 596, 211–220 (2021).

3. J. A. Cardin, M. C. Crair, M. J. Higley, Mesoscopic Imaging: Shining a Wide Light on Large-Scale Neural Dynamics. Neuron 108, 33–43 (2020).

4. Y. Xue, Q. Yang, G. Hu, K. Guo, L. Tian, L. Tian, L. Tian, Deep-learning-augmented computational miniature mesoscope. Optica 9, 1009–1021 (2022).

5. I. de Kernier, A. Ali-Cherif, N. Rongeat, O. Cioni, S. Morales, J. Savatier, S. Monneret, P. Blandin, Large field-of-view phase and fluorescence mesoscope with microscopic resolution. J Biomed Opt 24, 036501 (2019).

6. A. Glaser, J. Chandrashekar, J. Vasquez, C. Arshadi, N. Ouellette, X. Jiang, J. Baka, G. Kovacs, M. Woodard, S. Seshamani, K. Cao, N. Clack, A. Recknagel, A. Grim, P. Balaram, E. Turschak, A. Liddell, J. Rohde, A. Hellevik, K. Takasaki, L. E. Barner, M. Logsdon, C. Chronopoulos, S. de Vries, J. Ting, S. Perlmutter, B. Kalmbach, N. Dembrow, R. C. Reid, D. Feng, K. Svoboda, Expansion-assisted selective plane illumination microscopy for nanoscale imaging of centimeter-scale tissues. Elife 12 (2023).

7. A. E. Cohen, C. A. Werley, M.-P. Chien, Ultrawidefield microscope for high-speed fluorescence imaging and targeted optogenetic stimulation. Biomed Opt Express 8, 5794–5813 (2017).

8. T. Ichimura, T. Kakizuka, K. Horikawa, K. Seiriki, A. Kasai, H. Hashimoto, K. Fujita, T. M. Watanabe, T. Nagai, Exploring rare cellular activity in more than one million cells by a transscale scope. Sci Rep 11, 1–16 (2021).

9. T. Ichimura, T. Kakizuka, Y. Sato, K. Itano, K. Seiriki, H. Hashimoto, H. Itoga, S. Onami, T. Nagai, Volumetric trans-scale imaging of massive quantity of heterogeneous cell populations in centimeter-wide tissue and embryo. Elife 13 (2024).

10. R. Shi, X. Chen, J. Deng, J. Liang, K. Fan, F. Zhou, P. Tang, L. Zhang, L. Kong, Random-access wide-field mesoscopy for centimetre-scale imaging of biodynamics with subcellular resolution. Nat Photonics, 1–10 (2024).

11. G. McConnell, J. Trägårdh, R. Amor, J. Dempster, E. Reid, W. B. Amos, A novel optical microscope for imaging large embryos and tissue volumes with sub-cellular resolution throughout. Elife 5 (2016).

12. C. Zuo, J. Li, J. Sun, Y. Fan, J. Zhang, L. Lu, R. Zhang, B. Wang, L. Huang, Q. Chen, Transport of intensity equation: a tutorial. Opt Lasers Eng 135, 106187 (2020).

13. M. Mir, J. Rogers, H. Ding, S. Unarunotai, Z. Wang, L. Millet, M. U. Gillette, G. Popescu, Spatial light interference microscopy (SLIM). Opt Express 19, 1016–1026 (2011).

14. P. Marquet, C. Depeursinge, P. J. Magistretti, Review of quantitative phase-digital holographic microscopy: promising novel imaging technique to resolve neuronal network activity and identify cellular biomarkers of psychiatric disorders. Neurophotonics 1, 020901 (2014).

15. L. Durdevic, L. Durdevic, A. R. Ginés, A. Roueff, G. Blivet, G. Baffou, Biomass measurements of single neurites in vitro using optical wavefront microscopy. Biomed Opt Express 13, 6550–6560 (2022).

16. P. C. Chaumet, P. Bon, G. Maire, A. Sentenac, G. Baffou, Quantitative phase microscopies: accuracy comparison. ArXiv 2403 (2024).

17. M. M. Frigault, J. Lacoste, J. L. Swift, C. M. Brown, Live-cell microscopy – tips and tools. J Cell Sci 122, 753–767 (2009).

18. E. R. Polanco, T. E. Moustafa, A. Butterfield, S. D. Scherer, E. Cortes-Sanchez, T. Bodily, B. T. Spike, B. E. Welm, P. S. Bernard, T. A. Zangle, Multiparametric quantitative phase imaging for real-time, single cell, drug screening in breast cancer. Commun Biol 5, 1–12 (2022).

19. Y. Liu, S. Uttam, Perspective on quantitative phase imaging to improve precision cancer medicine. J Biomed Opt 29, S22705–1 (2024).

20. S. De Feraudy, I. Revet, V. Bezrookove, L. Feeney, J. E. Cleaver, A minority of foci or pan-nuclear apoptotic staining of γH2AX in the S phase after UV damage contain DNA double-strand breaks. Proc Natl Acad Sci U S A 107, 6870–6875 (2010).

21. C. J. Kim, A. L. Gonye, K. Truskowski, C. F. Lee, Y. K. Cho, R. H. Austin, K. J. Pienta, S. R. Amend, Nuclear morphology predicts cell survival to cisplatin chemotherapy. Neoplasia 42, 100906 (2023).

22. L. McInnes, J. Healy, N. Saul, L. Großberger, UMAP: Uniform Manifold Approximation and Projection. J Open Source Softw 3, 861 (2018).

23. X. Sun, P. D. Kaufman, Ki-67: more than a proliferation marker. Chromosoma 127, 175 (2018).

24. H. Was, A. Borkowska, A. Olszewska, A. Klemba, M. Marciniak, A. Synowiec, C. Kieda, Polyploidy formation in cancer cells: How a Trojan horse is born. Semin Cancer Biol 81, 24–36 (2022).

25. A. Klimas, B. R. Gallagher, P. Wijesekara, S. Fekir, E. F. DiBernardo, Z. Cheng, D. B. Stolz, F. Cambi, S. C. Watkins, S. L. Brody, A. Horani, A. L. Barth, C. I. Moore, X. Ren, Y. Zhao, Magnify is a universal molecular anchoring strategy for expansion microscopy. Nat Biotechnol 41, 858–869 (2023).

26. Gail McConnell, Where next for the Mesolens?, Wiley Analytical Science (2020). https://analyticalscience.wiley.com/content/article-do/next-mesolens.

27. G. Zheng, R. Horstmeyer, C. Yang, Wide-field, high-resolution Fourier ptychographic microscopy. Nature Photonics 7, 739–745 (2013).

28. H. Ma, R. Fu, J. Xu, Y. Liu, A simple and cost-effective setup for super-resolution localization microscopy. Sci Rep 7, 1–9 (2017).

29. H. Ma, Y. Liu, Super-resolution localization microscopy: Toward high throughput, high quality, and low cost. APL Photonics 5, 60902 (2020).

30. H. Ma, M. Chen, P. Nguyen, Y. Liu, Toward drift-free high-throughput nanoscopy through adaptive intersection maximization. Sci Adv 10, 7765 (2024).

31. Z. Bian, C. Guo, S. Jiang, J. Zhu, R. Wang, P. Song, Z. Zhang, K. Hoshino, G. Zheng, Autofocusing technologies for whole slide imaging and automated microscopy. J Biophotonics 13, e202000227 (2020).

32. H. Ma, Y. Liu, Super-Resolution Imaging through Single-Molecule Localization. Biomedical Optical Imaging, 4-1-4–26 (2021).

33. H. Ma, W. Jiang, J. Xu, Y. Liu, Enhanced super-resolution microscopy by extreme value based emitter recovery. Sci Rep 11, 1–10 (2021).

34. H. Ma, J. Xu, Y. Liu, WindSTORM: Robust online image processing for high-throughput nanoscopy. Sci Adv 5 (2019).

35. Connor Meehan, Jonathan Ebrahimian, Wayne Moore, Stephen Meehan, Uniform Manifold Approximation and Projection (UMAP), MATLAB Central File Exchange (2022). https://www.mathworks.com/matlabcentral/fileexchange/71902-uniform-manifold-approximation-and-projection-umap.

36. D. Paganin, A. Barty, A. Roberts, K. A. Nugent, Quantitative optical phase microscopy. Opt Lett 23, 817–819 (1998).

