## Supplementary Materials for "An Omni-Mesoscope for multiscale high-throughput quantitative phase imaging of cellular dynamics and high-content molecular characterization"

Hongqiang Ma *et al.*

**This PDF file includes:**

Supplementary Notes
Supplementary Figures S1 to S4
Supplementary Tables S1

### SUPPLEMENTARY FIGURES

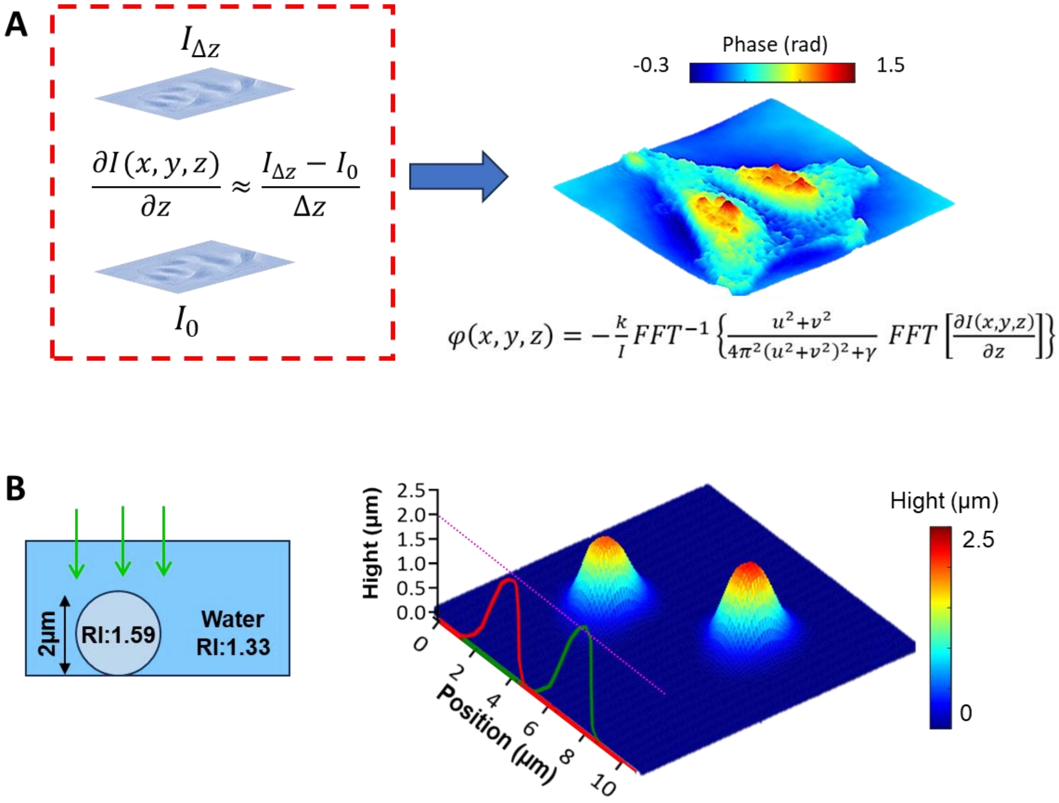


**Fig. S1**. **Evaluation of TIE-based quantitative phase imaging.** (**A**) Basis of TIE-based QPI is based on calculating the image gradient along the axial dimension. (B) Validation of the performance of TIE-based QPI using polystyrene microspheres with a diameter of ~2 µm.

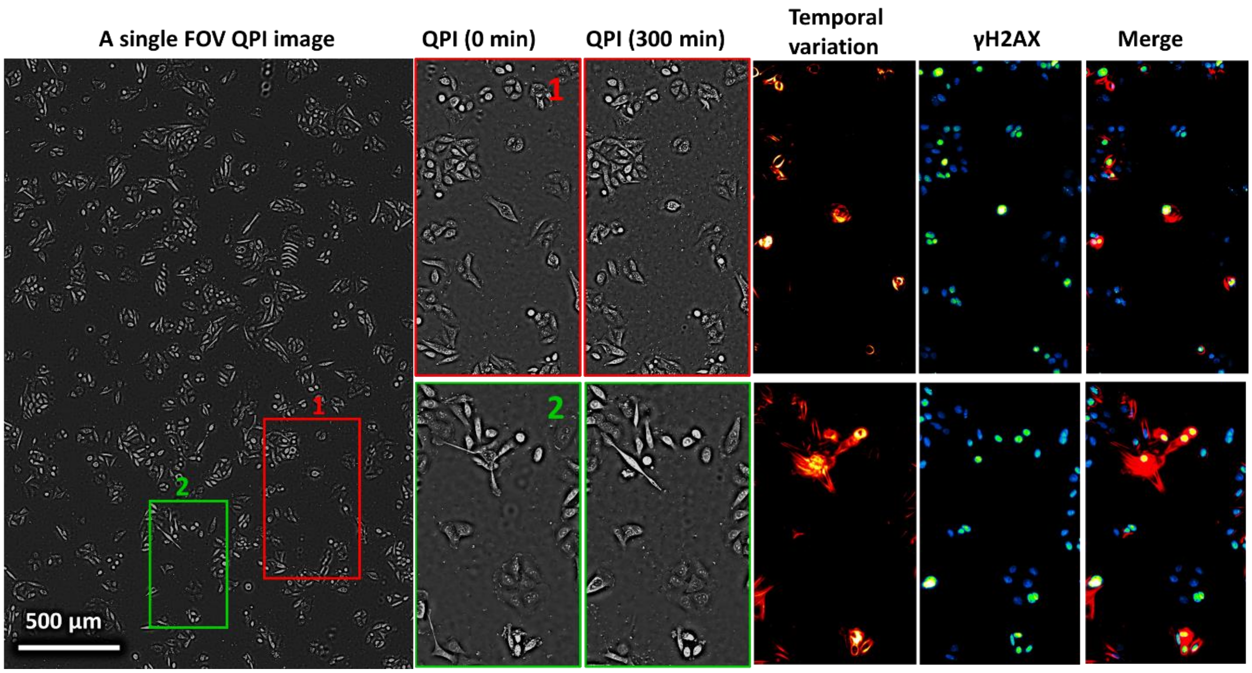


**Fig. S2**. **Correlation of dynamic cellular behaviors with DNA damage (γH2AX) in SW480 cells treated with 5-FU.** A representative quantitative phase image of SW480 cells treated with 35μM 5FU for 3 days is captured within a single field of view containing thousands of cells. Representative sequential snapshot images from the red and green boxes illustrate the dynamic morphological changes over 5 hours. We utilized the temporal standard deviation of the QPI videos to evaluate the local cell dynamics and movement. Most cells exhibit minimal movement, as indicated by a low temporal phase variation, while a subset demonstrates significant local motion, depicted by a high temporal phase variation. The cells displaying larger motions correspond to a higher DNA damage level, as evidenced by the higher intensity in the γH2AX image.

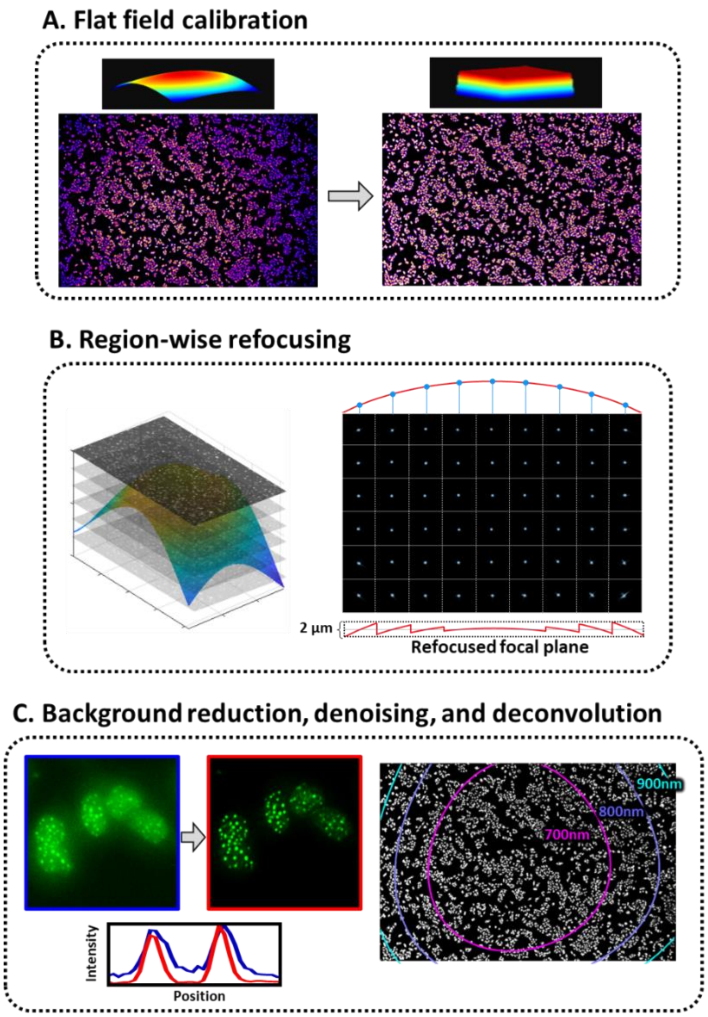


**Fig. S3**. **Three major steps to enhance the final image quality of Omni-Mesoscope.** (**A**) Flat-field calibration: Two strategies were employed to achieve uniform fluorescence imaging. (**B**) Region-wise refocusing: Given that all objective lenses exhibit field curvature, region-wise refocusing was implemented to address defocusing issues at the edge of the field of view as described in Methods. (**C**) Background reduction, denoising, and deconvolution.

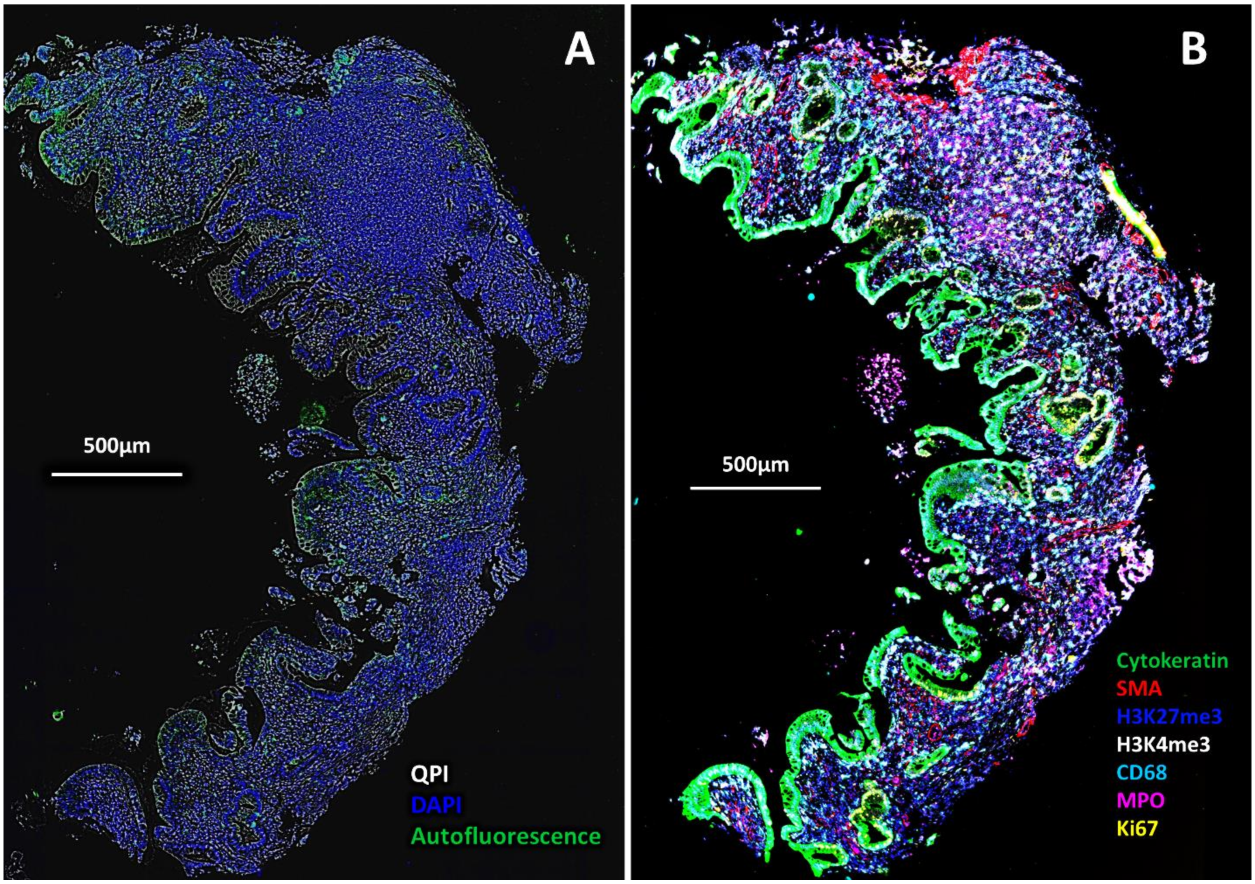


**Fig. S4**. **The multiplex fluorescence image of a human tissue section with ulcerative colitis, captured using our Omni-Mesoscope.** (**A**) This composite image incorporates a quantitative phase image, a DAPI fluorescence image, and an autofluorescence image excited by a 470nm laser. (**B**) The multiplex image showcases biomarkers including Cytokeratin, SMA, H3K27me3.H3K4me3, CD68, MPO, and Ki67. The image is generated by stitching together two single field of view (FOV) images with an overlapping ratio of 10%.

**Table S1**: Detailed information about the antibodies used for multiplex fluorescence imaging

| Antibody | Vendor | Catalog number | Dilutions | Cycle | Conjugated Dye |
|---|---|---|---|---|---|
| LMNAC | Cell Signaling Technology | 34698SF | 1:400 | 1 | None |
| H3K27me3 | Cell Signaling Technology | 9733S | 1:200 | 1 | None |
| Goat Anti-Rabbit IgG (H+L) | Jackson ImmunoResearch | 111-005-144 | 1:200 | 1 | CF568 |
| Goat Anti-Mouse IgG (H+L) | Jackson ImmunoResearch | 115-005-166 | 1:100 | 1 | AlexaFluor 647 |
| H3 | Santa Cruz Biotech | sc-517576 | 1:100 | 2 | AlexaFluor 405 |
| Ki67 | Cell Signaling Technology | 9449S | 1:400 | 2 | Dy488 |
| Tubulin | Cell Signaling Technology | 5059S | 1:400 | 2 | AlexaFluor 555 |
| MCM7 | Santa Cruz Biotech | sc-9966 | 1:50 | 2 | AlexaFluor 594 |
| CENP-A | Thermo Fisher Scientific | MA1-20832 | 1:50 | 2 | CF647 |
| P53 | Cell Signaling Technology | 5429S | 1:100 | 3 | AlexaFluor 488 |
| Vimentin | BioLegend | 677804 | 1:200 | 3 | AlexaFluor 555 |
| H2Ax | Millipore Sigma | 05-636 | 1:100 | 3 | AlexaFluor 594 |
| H3K9me3 | Abcam | ab8898 | 1:400 | 3 | AlexaFluor 647 |
| HP1 | Santa Cruz Biotech | sc-515341 | 1:100 | 4 | AlexaFluor 405 |
| EZH2 | Abcam | ab283294 | 1:100 | 4 | Dy488 |
| P21 | Cell Signaling Technology | 8493S | 1:100 | 4 | AlexaFluor 555 |
| CD44 | Biolegend | 103054 | 1:100 | 4 | AlexaFluor 594 |
| H3K9ac | Abcam | ab12179 | 1:400 | 4 | AlexaFluor 647 |
| H3K27ac | Abcam | ab177178 | 1:200 | 5 | AlexaFluor 405 |
| Cytokeratin | Invitrogen | 53-9003-82 | 1:200 | 5 | AlexaFluor 488 |
| H3K4me3 | Abcam | ab8580 | 1:200 | 5 | CF568 |
| RNAPII | Abcam | ab5408 | 1:200 | 5 | AlexaFluor 657 |
